# A Compact Quadrupole-Orbitrap Mass Spectrometer with FAIMS Interface Improves Proteome Coverage in Short LC Gradients

**DOI:** 10.1101/860643

**Authors:** Dorte B. Bekker-Jensen, Ana Martínez del Val, Sophia Steigerwald, Patrick Rüther, Kyle Fort, Tabiwang N. Arrey, Alexander Harder, Alexander Makarov, Jesper V. Olsen

**Affiliations:** The Novo Nordisk Foundation Center for Protein Research, University of Copenhagen, DENMARK; Thermo Fisher Scientific, Bremen, GERMANY

**Keywords:** Bottom-up Shotgun proteomics Orbitrap DIA FAIMS DDA phosphoproteomics TMT

## Abstract

State-of-the-art proteomics-grade mass spectrometers can measure peptide precursors and their fragments with ppm mass accuracy at sequencing speeds of tens of peptides per second with attomolar sensitivity. Here we describe a compact and robust quadrupole-orbitrap mass spectrometer equipped with a front-end High Field Asymmetric Waveform Ion Mobility Spectrometry (FAIMS) Interface. The performance of the Orbitrap Exploris 480 mass spectrometer is evaluated in data-dependent acquisition (DDA) and data-independent acquisition (DIA) modes in combination with FAIMS. We demonstrate that different compensation voltages (CVs) for FAIMS are optimal for DDA and DIA, respectively. Combining DIA with FAIMS using single CVs, the instrument surpasses 2500 unique peptides identified per minute. This enables quantification of >5000 proteins with short online LC gradients delivered by the Evosep One LC system allowing acquisition of 60 samples per day. The raw sensitivity of the instrument is evaluated by analyzing 5 ng of a HeLa digest from which >1000 proteins were reproducibly identified with 5 minute LC gradients using DIA-FAIMS. To demonstrate the versatility of the instrument we recorded an organ-wide map of proteome expression across 12 rat tissues quantified by tandem mass tags and label-free quantification using DIA with FAIMS to a depth of >10,000 proteins.

## INTRODUCTION

Deep proteome profiling of human cells can now be routinely achieved by shotgun proteomics using the latest advances in nanoflow liquid chromatography tandem mass spectrometry (LC-MS/MS)^1,2^. However, comprehensive analysis of a human proteome typically requires days of mass spectrometric measurement time^3^, making high-throughput analysis challenging and time-consuming. Therefore, new improved mass spectrometric technology and acquisition strategies are needed to overcome these limitations in current large-scale proteomics. The Orbitrap analyzer has become one of the major players in mass spectrometry-based proteomics^4^. We have previously described the performance enhancement achieved over the different generations of the popular benchtop quadrupole-Orbitrap instruments, the Q Exactive™ series, for proteomics applications^5–7^. The previous generation, the Q Exactive HF-X™ hybrid quadrupole-Orbitrap instrument, incorporated a new peak-picking algorithm, a brighter ion source, and optimized ion transfers enabling productive HCD-MS/MS^8^ acquisition above 40 Hz. These improvements collectively resulted in increased peptide and protein identification rates of up to fifty percent at short LC-MS gradients with more than 1000 unique peptides identified per minute^7^. However, instrument performance after extended periods of continuous sample analyzes could be comprised by contaminants entering the quadrupole and resulting in local charging and lower MS/MS sensitivity. This meant that regular preventative maintenance could be necessary to preserve high performance of the Q Exactive HF-X™ instrument. Moreover, the semi-stochastic nature of the data-dependent acquisition (DDA) - the most popular tandem mass spectrometric acquisition strategy may create issues with reproducibility of peptide measurements between samples leading to the so-called missing value problem. To overcome the precursor selection problem inherent to DDA, data-independent acquisition (DIA) offers systematic measurement of all peptide ions regardless of their intensity by co-fragmenting all co-eluting peptides in broader precursor isolation windows. This provides wider dynamic range of the proteomes analyzed, improved reproducibility for identification and enabled better sensitivity and accuracy for quantification. The highly multiplexed fragment ion spectra in DIA require more elaborate processing algorithms for identification and quantification. These typically rely on spectral libraries previously recorded by data-dependent acquisition of similar sample types.

The front-end high-field asymmetric waveform ion mobility spectrometry (FAIMS)^9–11^ interface functions as an ion selection device and an electrospray filter that prevents neutrals from entering the orifice of the mass spectrometer while reducing chemical background noise. This ‘purification’ of the electrosprayed ions typically results in improved robustness and sensitivity for proteomics experiments. The FAIMS system continuously selects and focuses ions at atmospheric pressure based on their differential mobility in a high field vs. a low electric field established by applying an asymmetric (e.g., bi-sinusoidal) voltage to at least one of the inner and outer electrodes. To prevent an ion species from being discharged, the FAIMS system applies a small DC potential, called the compensation voltage (CV), to the inner electrode to compensate for the ion drift resulting from the applied asymmetric waveform. Selection of an ion species is based on a combination of its gas-phase charge-state and collisional cross-section. CVs can be tuned such that only selected subsets of ions are transmitted through the electrodes thereby favoring multiply-charged peptide species and filtering out singly-charged chemical background ions. The CV is typically negative for positive ions and positive for negative ions. The advantage of FAIMS in the context of large-scale proteomics on the Tribrid™ Orbitrap systems (i.e., those incorporating a quadrupole mass filter and dual-pressure linear quadrupole ion trap in addition to the Orbitrap mass analyzer) has previously demonstrated to improve proteome coverage for single-shot proteomics with long gradients by CV sweeping^11^ and reduce ratio compression in TMT experiment^12–14^. However, potential advantages of FAIMS for short gradients using single CVs have not been exploited. Likewise, FAIMS in combination with DIA has not yet been investigated. Here we describe a thorough performance evaluation of the hybrid quadrupole-Orbitrap Exploris 480 MS in combination with short LC gradients using the Evosep One system. Standard DDA in combination with FAIMS identifies more proteins but fewer peptides resulting in a compromise between sequence coverage and proteome depth. This is not the case for DIA, where FAIMS results in identification of more proteins while maintaining the number of peptide identifications.

## RESULTS AND DISCUSSION

### A compact quadrupole-Orbitrap instrument for proteomics

The Orbitrap Exploris 480 mass spectrometer combines a compact and highly integrated hybrid quadrupole-Orbitrap design with increased performance in high-throughput analysis. Compared to the Q Exactive HF™ ^15^ and HF-X™ ^7^ mass spectrometers that use the same combination of analyzers, the entire layout of the instrument was completely reconfigured to enable a more compact and robust platform, easier to service. At the same time, the new mass spectrometer shares a lot with the latest Tribrid instruments^16^ such as the Orbitrap Fusion Lumos™ mass spectrometer and Orbitrap Eclipse™ Tribrid™ mass spectrometer. The Orbitrap Exploris™ 480 instrument has the same ion source, atmosphere-to-vacuum interface and user-facing software as the Orbitrap Fusion Lumos™ mass spectrometer. The shared design of the Orbitrap Exploris 480 mass spectrometer also enables it to be interfaced to the same front-end options that are available with the Tribrid instruments, such as the FAIMS device and internal calibrant (EASY-IC™) source. Beyond this front end, everything else in the Orbitrap Exploris 480 instrument including the bent flatapole, quadrupole, independent charge detector and ion routing multipole was re-designed to rise to enhanced requirements of the instrument (Figure 1A). However, the backbone of the new design is a new pumping concept, combined with a new layout of ion optics, C-trap electrodes, and Orbitrap electrodes. The requirement of ultra-high vacuum (UHV) in the 10E-10 or even 10E-11 mbar range for Orbitrap analysis of ions formed at atmospheric pressure has previously necessitated a bulky, multi-stage pumping system, which represented the major constraint on the overall design and size of the instrument. The breakthrough realized in this new instrument concept is a single, purpose-developed, six-stage turbo pump that evacuates the entire path from the ion funnel to the Orbitrap analyzer. This enables a drastic reduction of instrument footprint and volume and enables easier access to all ion-optical components. Based on improved understanding of ion dynamics, both ion optics and instrument control software were extensively refined to reduce spectral noise and the influence of ion optics contamination. The latter is achieved by using accelerating-only metal lenses, symmetrical ion loading of the quadrupole mass filter and electrodynamic squeezing during ion transfer into the C-trap. Asymmetric ion extraction from the C-trap provides higher resistance to space charge-induced ion losses, while symmetric suspension of the Orbitrap electrodes reduces noise coupling and improves pumping. Compared to the Q Exactive HF-X™ mass spectrometer, the new instrument features higher maximum resolving power up to 480,000 at m/z 200, as well as offering compatibility with an optional internal calibration source and the FAIMS interface.

**Figure 1.**
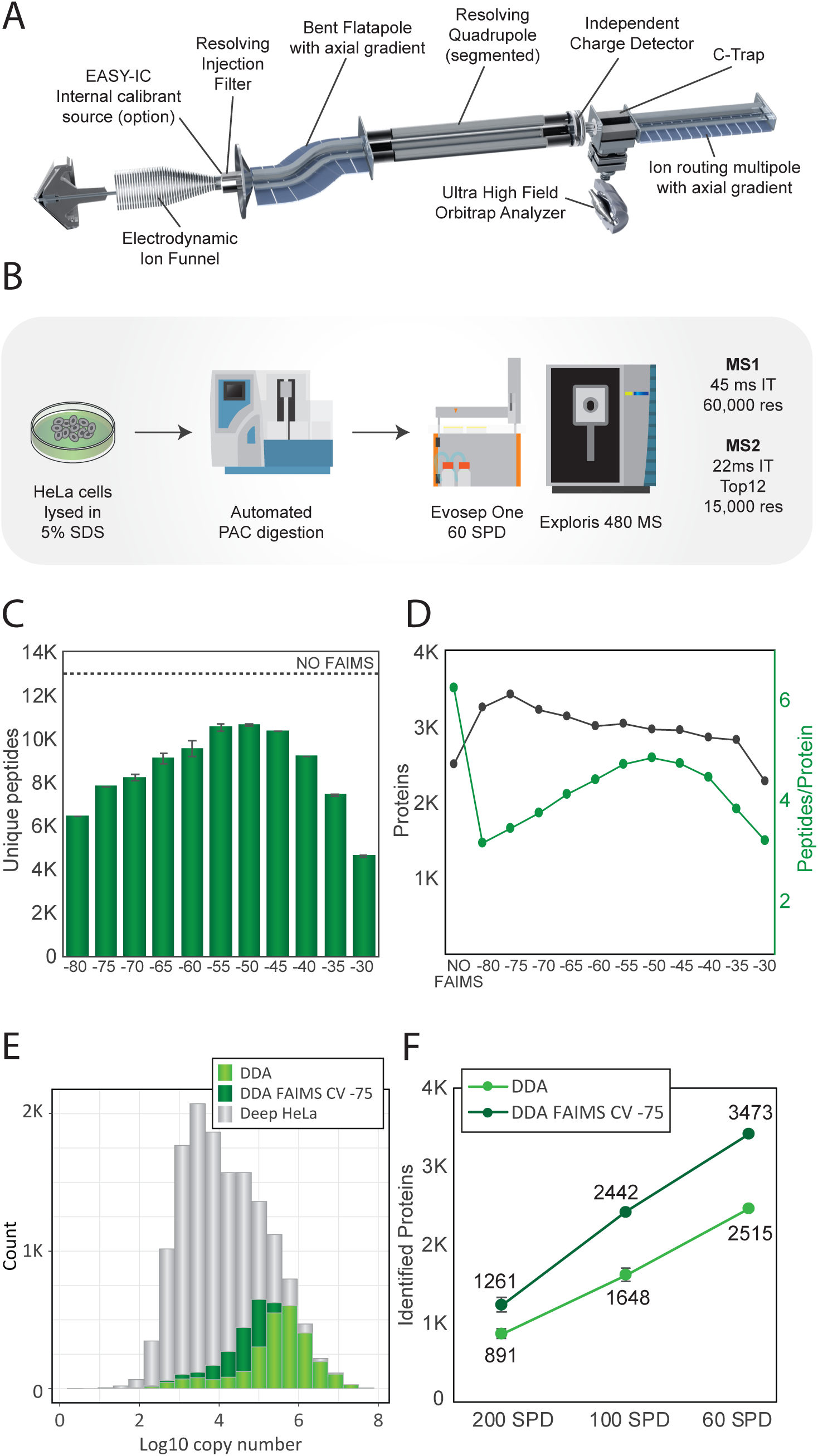
**(a)** Hardware overview of the Orbitrap Exploris 480 MS instrument. **(b)** Experimental workflow for HeLa measurements. **(c)** Barplot depicting the number of unique peptides identified using different compensation voltages with FAIMS. The number represented is the average of two replicates. Dashed line represents the average number of peptides identified in a DDA experiment without FAIMS. **(d)** Average number of proteins in black with and without using FAIMS with different compensation voltages. In green, number of peptides identified per protein. The plotted values are the average of at least two technical replicates. **(e)** Histogram of the protein abundance distribution represented as copy number in log10 scale of a deep HeLa proteome (gray) overlayed with 21 minutes single shot analyzes with FAIMS CV of -75 V (dark green) and without FAIMS (light green). **(f)** Number of proteins identified from 500 ng of peptide using DDA (light green) or DDA with FAIMS and optimal CV of -75 V (dark green) in different gradient lengths: 200 SPD, 100 SPD and 60 SPD.

### Data-dependent acquisition with FAIMS

To evaluate the performance of the instrument for proteomics we analyzed single-shot HeLa proteomes using short LC gradients both with and without FAIMS. HeLa cervix carcinoma cells were harvested in 5% SDS buffer and tryptic digests were prepared using an automated workflow by protein aggregation capture (PAC)^17^ and proteolytic digestion with endoproteinase Lys-C and trypsin on magnetic beads using a KingFisher™ Flex robot. To minimize overhead time between LC-MS/MS runs and maintain robustness and reproducibility of LC separations over hundreds of injections, we made use of the Evosep One^18^ system with short gradients in the range from 5 to 21 minutes enabling analysis of 60-200 samples per day. Although it utilizes a fundamentally different design, the Orbitrap Exploris 480 MS can achieve the same ms/ms scanning speed as the Q Exactive HF-X™ instrument when using the same transient and maximum injection times for optimal parallel mode of acquisition^7^. All data-dependent acquisition (DDA) experiments were therefore performed with a robust and fast scanning top12 method encompassing a MS1 survey scan recorded at 60,000 resolution followed by twelve HCD scans with maximum allowed injection time of 22 ms and 15,000 resolution equivalent of 28 Hz MS/MS scan speed (Figure 1B). To identify the optimal FAIMS settings for maximum proteome coverage we performed a systematic step-wise CV sweep from -80 V to -30 V with 5 V intervals and benchmarked the number of peptides and proteins identified at each CV setting with a standard DDA run without FAIMS. With 500 nanograms of HeLa peptides injected on column and analyzed with the 21 minutes (60 samples per day) gradient we observed a similar peptide distribution across the CV range as shown previously^11^ with maximum peptide identification for CV of -50 V (Figure 1C and Supplementary table 1). All DDA files were analyzed with MaxQuant^19^ applying a one percent false discovery rate (FDR) at both peptide and protein level. The reproducibility and separation power of the FAIMS device judged by the replica runs suggests that a difference of more than 10 V between CVs is required to minimize overlaps (Supplementary Figure 1A). At a CV of -40 V and lower, singly charged precursors were significantly reduced, whereas multi-charged precursors are spread across the CV range (Supplementary Figure 1B). None of the FAIMS files resulted in the same number of identifications as the run without FAIMS at 12,700 peptides. Conversely, all FAIMS runs except the one with a CV of -30 V achieved significantly higher number of protein identifications with a CV of -75 V providing the best coverage with 3473 proteins (representing 38% more than the 2515 proteins identified without FAIMS). However, the increased proteome depth with FAIMS comes at a cost of lower sequence coverage at single CVs (Figure 1D). Surprisingly, a CV of -75 V provides the best protein coverage, but the smallest number of peptides resulting in only about three peptides per protein on average (Figure 1D Supplementary table 1). At a CV of -75 V, we are operating at the edge of the peptide intensity distribution where a very small fraction of peptides is transmitted through the FAIMS device. The success of this CV setting is therefore highly dependent on the general performance of the instrument, as any loss in sensitivity will immediately result in a dramatic decrease in protein identification. It only works well when the total ion flux in MS/MS mode is high as measured by total ion chromatogram (TIC) distributions plot (Supplementary Figure 1C). To assess the dynamic range of the proteins covered we overlapped the identifications with the deepest HeLa proteome to date with copy number estimates for 12,600 protein-coding genes^3^. In the DDA-FAIMS experiments, we not only cover more proteins compared to the DDA without FAIMS, but we are also able to dig deeper into the HeLa protein abundance distribution covering significantly more of the dynamic range (Figure 1E). The observed increase in the number of proteins identified with FAIMS operated with a CV of -75 V compared to runs without FAIMS is true for all gradients tested with even higher gains at shorter gradients, possibly due to concentration of the MS/MS signal intensity (Figure 1F). Using the 200 samples per day gradient, we observe a 41% increase of proteins and 48% gain when using the 100 samples per day gradient (Figure 1F). Previously, it has been shown that in longer gradients switching CVs internally during the LC-MS/MS analysis improves peptide coverage^11^. However, with a current overhead time of 25 ms between individual single CVs, it is more suitable for gradients longer than the ones used in the current study.

### Data-independent acquisition with FAIMS

Next, we wanted to test if the benefits of applying FAIMS with a single CV could be extended to data-independent acquisition (DIA). We again started by performing a CV sweep from -80 V to - 35 V in steps of 5 V and compared the number of quantified peptides at each CV with an equivalent LC-MS/MS run without FAIMS. All DIA files were analyzed using the 60 samples per day gradient and processed with Spectronaut^20^ software using a project-specific spectral library recorded at the same chromatography settings consisting of ∼105,000 tryptic HeLa peptides and nearly 10,000 proteins for the analysis. We previously demonstrated that DIA provides higher proteome coverage than DDA with short gradients on a Q Exactive HF-X™ instrument^7^. This is also the case with the Orbitrap Exploris 480 MS with which we identified up to 4587 HeLa proteins in a single run using DIA or 35% more proteins than by DDA. Importantly, in contrast to the FAIMS-DDA analysis, we found that FAIMS-DIA worked best at a CV of -45 V with more identifications of peptides and 5138 proteins compared to a DIA run without FAIMS (Figure 2A and Supplementary table 2). Furthermore, DIA-FAIMS reach impressive identification rates of up to 2500 peptides per minute of gradient time (Figure 2B). This translates to a consistent gain in protein identifications across the gradient for DIA-FAIMS compared to classic DIA (Figure 2C). These results highlights how the use of FAIMS in DIA mode, increased proteome coverage but maintains the peptide identification rate in DIA mode. Even though the differential ion mobility separation by FAIMS results in a reduction in measured fragment ions, the increase in proteins identified is due to more ions are sampled in MS/MS mode in DIA-FAIMS compared to DDA (Supplementary Figure 2A-B). In proteomics, it is not only central to identify as many peptides and proteins as possible but high quantitative precision is equally important. To test the impact of FAIMS on the quantitative accuracy, we analyzed the reproducibility of protein intensities between replicates. For DIA-FAIMS 71% of the 5138 identified proteins were reproducibly quantified with a coefficient of variation below 20% and 54% with a coefficient of variation below 10% (Figure 2D). In contrast, DIA analyzes without FAIMS decreased the number of consistently quantified proteins with a coefficient of variation below 20% to 2829 representing only 60% of the total 4584 proteins quantified (Figure 2D). Importantly, this highlights that no compromise on quantitative accuracy is observed for the additional proteins identified with DIA-FAIMS. Compared with DDA-FAIMS, we observe that around fifty percent additional proteins identified in DIA-FAIMS match the same proteome abundance distribution range (Figure 2E).

**Figure 2.**
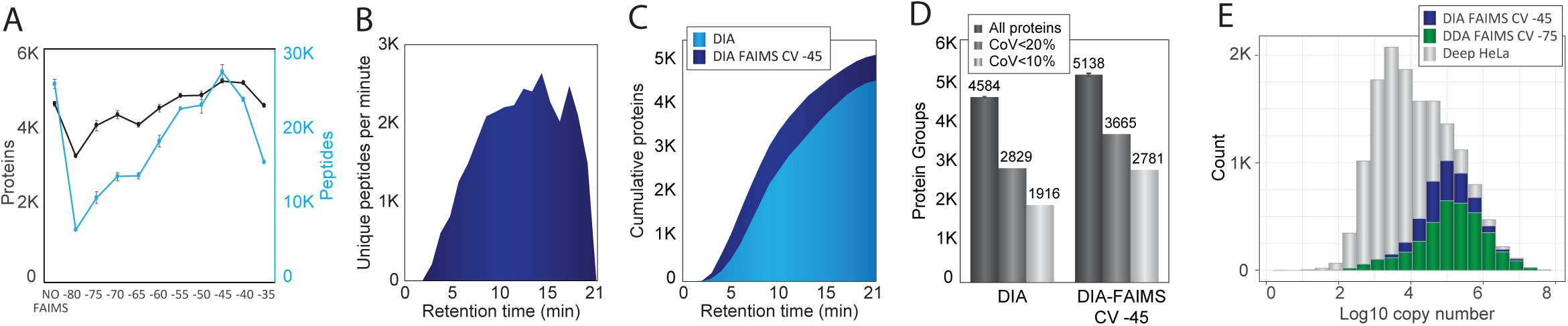
**(a)** In black, number of proteins quantified from 500 ng of peptide in 21 minutes in a DIA experiment without FAIMS and with FAIMS using different compensation voltages. In blue, number of peptides quantified in the same experiments. The plotted values are the average of at least two technical replicates. **(b)** Visualization of unique peptide identification across the gradient length for DIA run with CV of 45 V with FAIMS and 500ng load. **(c)** Cumulative protein identifications across the gradient for DIA runs without (light blue) and with CV of -45 V with FAIMS (dark blue)**. (d)** Bar chart of all protein identifications (dark grey), proteins with coefficient of variation below 20% (grey) and below 10% (light grey) for DIA with and without CV of -45 V with FAIMS. **(e)** Histogram of the protein abundance distribution represented as copy number in log10 scale of a deep HeLa proteome (gray), and in a single shot analysis of 21 minutes using DDA and optimal CV of -75 V with FAIMS (dark green) or using DIA and optimal CV of -45 V with FAIMS (blue).

### Sensitive analysis with FAIMS and very short gradients

Given the significant increase in protein identifications with 500 ng HeLa digest analyzed with FAIMS using the 60 samples per day gradient, we wondered whether this gain could also be achieved at decreased sample load. For optimal sensitivity, the gradient was shortened to 5 min (200 samples per day). We analyzed 5 ng of tryptic peptides loaded on column equivalent to approximately 30-50 HeLa cells. The samples were analyzed with the short 5 min gradient (200 samples per day) using both DDA and DIA with and without FAIMS. For such low amounts, it is required to enhance the MS/MS sensitivity by increasing the resolving power and injection times to 30,000 resolution and 54 ms, respectively. Moreover, we observed that a slightly shifted CV to-50 V is optimal (Supplementary Figure 3A). To maintain good quantitative accuracy it is required to reduce the cycle time from 2 s to 1 s (Supplementary Figure 3B). The very low amount is a challenge for standard DDA, where we only identify 26 proteins. This is a setting where FAIMS is valuable by increasing protein identification by a factor of 10 to 281 proteins. On the other hand, classic DIA further increased proteins identified to 547, while DIA-FAIMS is superior with more than 1000 proteins identified (Figure 3A Supplementary table 3). Additionally, FAIMS enhanced the reproducibility of DIA quantification, based on the number of proteins quantified with coefficient of variation below 20% (Figure 3B). The observed ten-fold increase in number of proteins for the DDA-FAIMS compared to DDA without FAIMS suggests that DDA-FAIMS is ideally suited for DDA analysis of very low input material. The main reason for this is that FAIMS is capable of effectively removing the singly-charged background ions from ambient air and the electrospray buffer, which otherwise take up a lot of C-trap capacity when minute amounts of peptide mixtures are analyzed (Figure 5E). This issue is best illustrated by the scaled total ion chromatograms (TICs), where the background level in the DDA experiment is very high (Figure 3C), whereas with DDA-FAIMS the background is very low and hence the peptide signals are significantly higher (Figure 3D). This enhancement in the spectral quality by FAIMS is also striking when quantifying the number of ions measured in the full-scan MS1 spectrum during the gradient, where FAIMS has a much lower starting background level but ten-fold higher levels during peptide elution from 1.5 to 4 minutes (Figure 3F).

**Figure 3.**
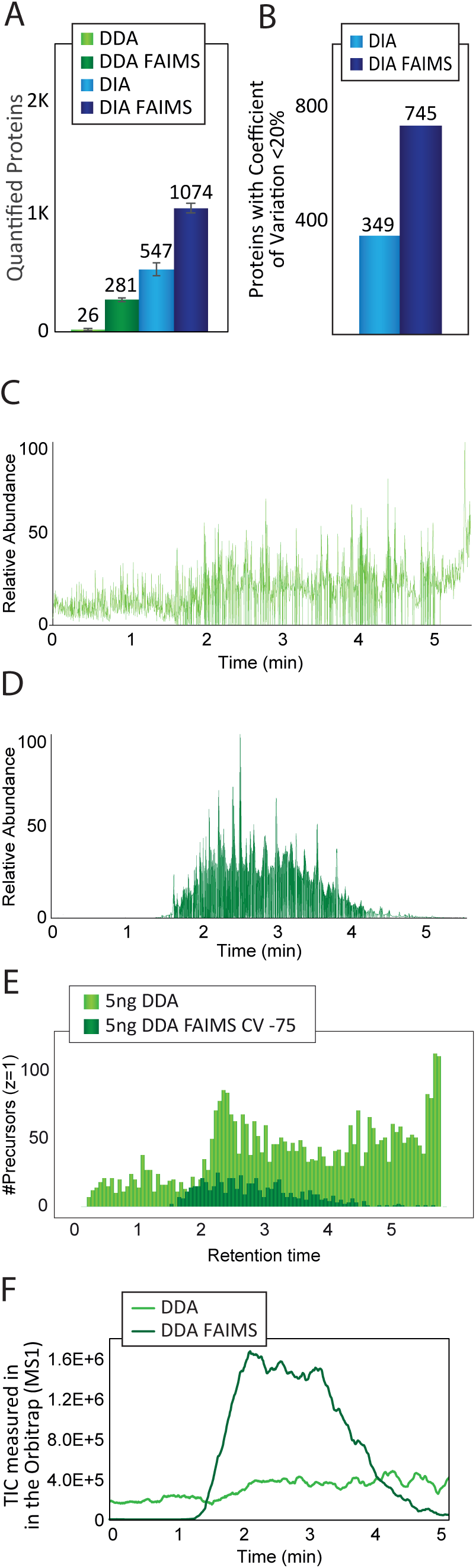
**(a)** Bar chart showing the number of proteins identified in 5 min gradients (200 samples per day) using 5 ng load. Light green bars correspond to DDA acquisition settings, dark green bars correspond to DDA with optimal CV with FAIMS, light blue bars correspond to DIA acquisition and dark blue bars correspond to DIA using FAIMS. Plotted values are the average between at least three replicates. **(b)** Bar chart showing the number of proteins quantified with a coefficient of variation below 20% in DIA experiments with FAIMS (dark blue) or without FAIMS (light blue). **(c)** Chromatogram of a DDA run from 5 ng using a 5 minutes gradient showing the relative abundance of the total ion current signal. **(d)** Chromatogram of a DDA run with FAIMS (CV -75) from 5 ng using a 5 minutes gradient showing the relative abundance of the total ion current signal. **(e)** Histogram showing the abundance distribution of precursors with charge +1 during a 5 minutes gradient using a sample loading of 5ng in a DDA run (light green) or a DDA with FAIMS at CV -75 (dark green). **(f)** Plot of the TIC measured in the Orbitrap analyzer in MS1 scans across time in a 5 minute gradient using a sample loading of 5ng in a DDA run (light green) or a DDA with FAIMS at CV-75 (dark green).

**Figure 4.**
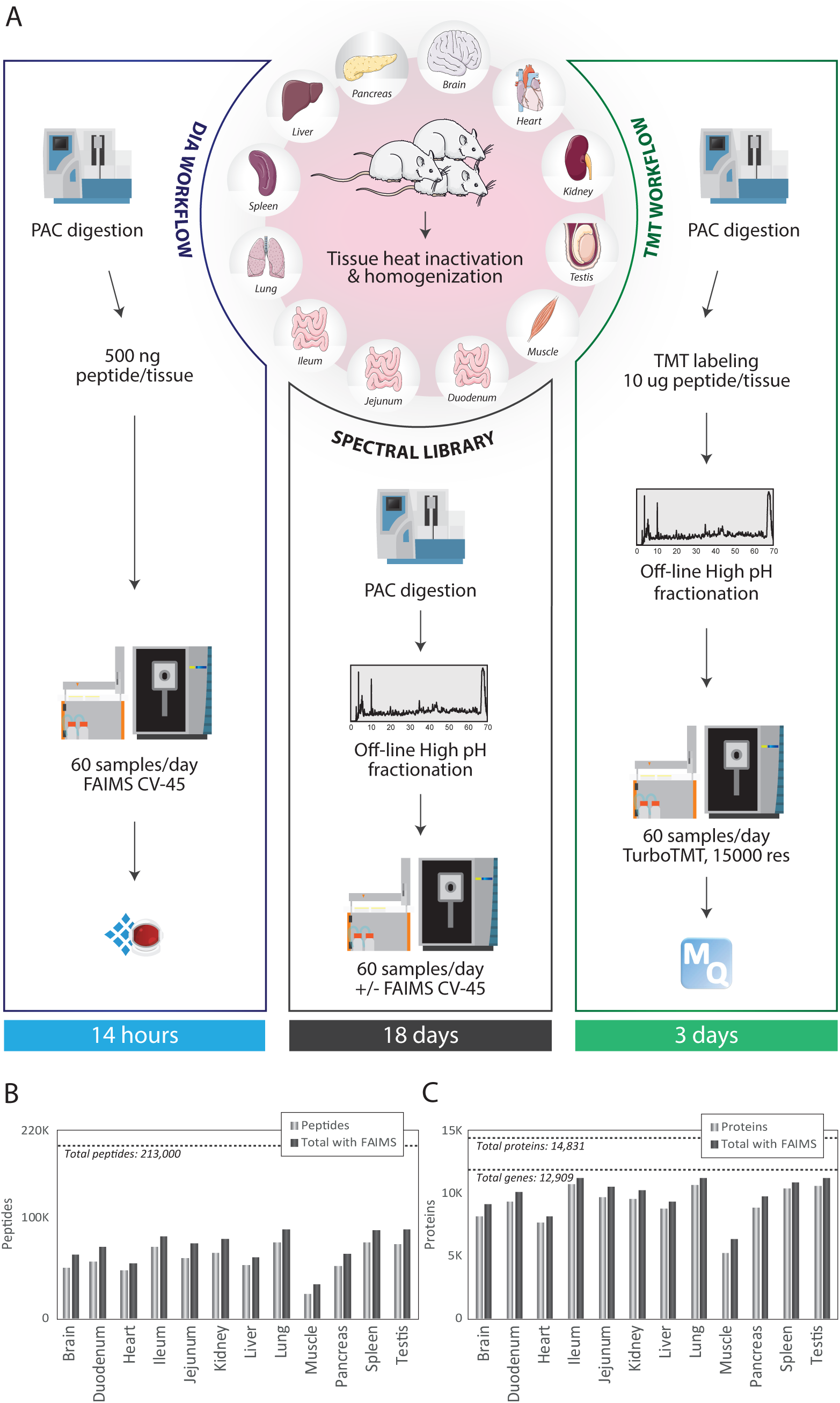
**(a)** Experimental workflow of the rat tissue atlas. **(b)** Bar chart showing the number of peptides quantified in each single tissue library (light gray bar) and the gain in peptides after merging two libraries for the same tissue, one acquired without FAIMS and one acquired with FAIMS at CV-45 (dark gray). Dashed line indicates the total number of peptides identified in the complete library generated from the 12 tissues acquired with and without FAIMS. **(c)** Bar chart showing the number of protein groups quantified in each single tissue library (light gray bar) and the gain in peptides after merging two libraries for the same tissue, one acquired without FAIMS and one acquired with FAIMS at CV-45 (dark gray). Dashed lines indicate the total number of protein groups and protein coding genes quantified in the complete library generated from the 12 tissues acquired with and without FAIMS.

**Figure 5.**
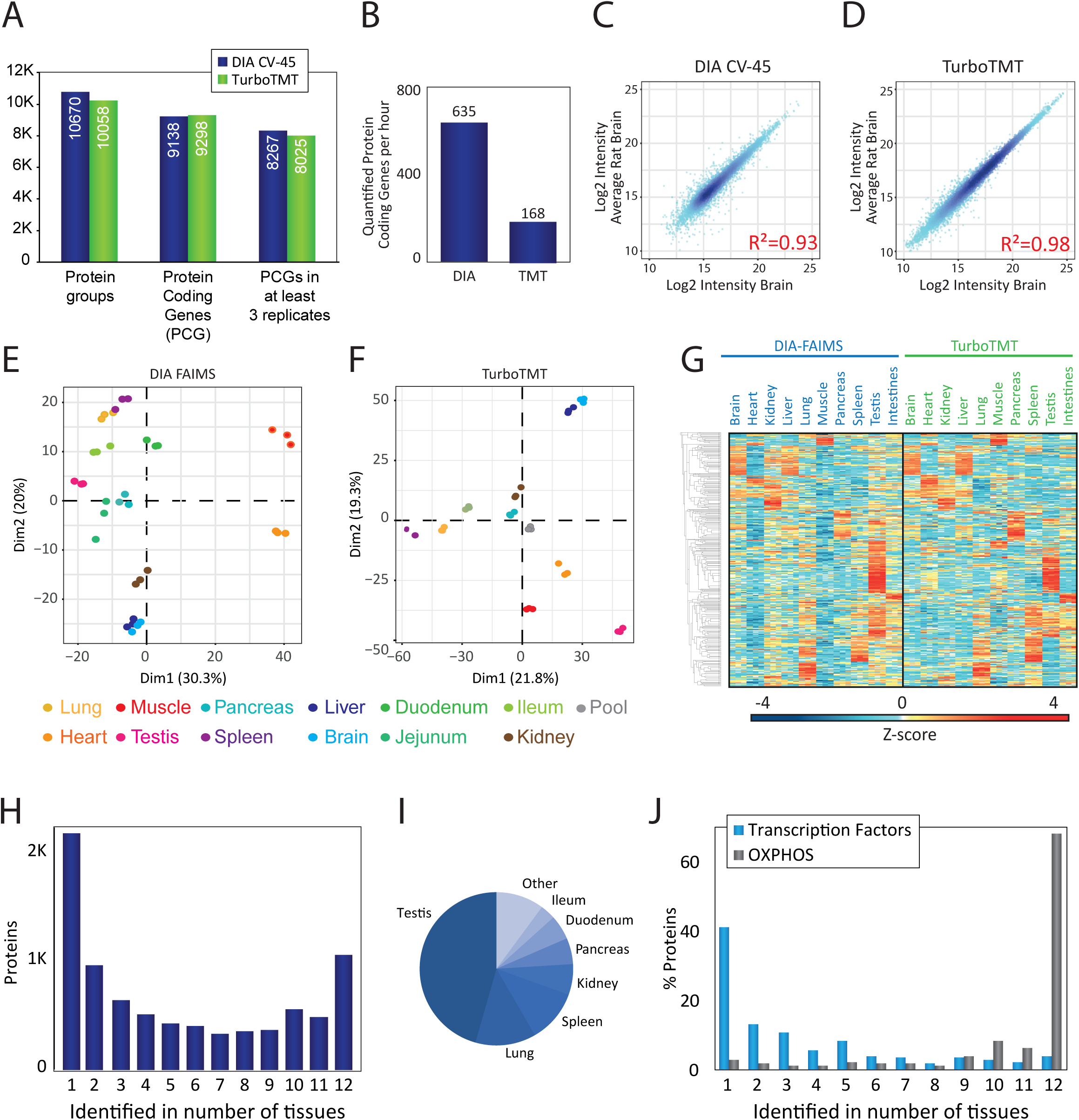
**(a)** Bar chart showing the number of protein groups, protein coding genes (PCGs) identified and PCGs quantified in three replicates of the same tissue using DIA and FAIMS (dark blue) or turboTMT (green). **(b)** Number of PCGs per hour quantified in the DIA-FAIMS and turboTMT experiments. **(c)** Correlation plot of the log2 intensity of each protein quantified in each rat brain against the average of the other two replicates from the same tissue in DIA-FAIMS. Correlation is measured as R-squared. **(d)** Correlation plot of the log2 intensity of each protein quantified in each rat brain against the average of the other two replicates from the same tissue in turbo-TMT. Correlation is measured as R-squared. **(e)** Principal Component Analysis (PCA) showing the classification of the 12 tissues from the three rats analyzed in the DIA-FAIMS experiment. **(f)** Principal Component Analysis (PCA) showing the classification of the 10 tissues and the pooled sample from the three rats analyzed in the turbo-TMT experiment. **(g)** Hierarchical clustering of the intensity (plotted as z-score) of the proteins identified in DIA-FAIMS and turbo-TMT experiments. **(h)** Bar chart describing the number of specific proteins identified across tissues, where 1 indicates that the protein has only been identified in three replicates of the same tissue and 12 indicates that it has been quantified in all tissues. **(i)** Pie chart showing the tissue-specificity of proteins identified in only one tissue. **(j)** Bar chart depicting the percentage of proteins, either transcription factors (blue) or enzymes from the Oxidative Phosphorylation pathway (gray), expressed in different number of tissues.

### Rat tissue specific protein atlas

The ability to quantify proteins and their post-translational modifications between cell states or cell types at a large scale is important for understanding protein regulation and dynamics in health and disease. The Orbitrap Exploris 480 mass spectrometer with its fast sequencing speed and sensitivity is ideally suited for deep proteome profiling across many conditions. To identify the optimal quantitative proteomics workflow for large-scale proteome expression studies, we compared tandem mass tags (TMT11-plex) with DIA-based label-free quantification of 12 tissues covering the major mammalian organs from three individual male rats (Figure 4A). To preserve the in-vivo state of the organ proteomes and phosphoproteomes, the excised tissues were heat-inactivated^21^ and homogenized in 5% SDS buffer. Extracted proteins were PAC precipitated on magnetic beads and digested with Lys-C and trypsin. All LC-MS/MS experiments were analyzed using the 21 min gradient (60 samples per day) on the Evosep One. For the DIA-based label-free quantification we analyzed each sample with the application of FAIMS to evaluate if it also provides benefits for complex organ proteome analyzes. Since spectral libraries are required for DIA, we performed deep proteome profiling by offline high-pH (HpH) reversed phase fractionation of the organ digests and analyzed each fraction by DDA analysis with and without FAIMS. Analyzing 46 HpH fractions from each of the 12 tissues +/-FAIMS at CV of -45 V resulted in 1104 LC-MS/MS runs. While each tissue required 32 hours of LC-MS measurement time, all tissues collectively needed 18.4 days of total mass spectrometric measurement time. The peptide coverage in each tissue varied from 50,000 unique peptides identified in skeletal muscle to 100,000 identified in testis (Figure 4B). The number of identified peptides generally correlates with the number of proteins identified in each organ with testis representing more than ten thousand proteins (Figure 4C). Combining the DDA analyzes with and without FAIMS slightly improves the depth of the library on both peptide and protein level. The observed discrepancy in identifications likely reflects the differences in protein expression levels across different organ proteomes, where highly abundant contractile proteins such as myosin dramatically increases the dynamic range of the expressed muscle proteome^22^. This high-quality dataset representing 213,000 unique peptides covering 12,909 protein-coding genes should represent a resource for the proteomics community, which can be bioinformatically mined or used as a general rodent spectral library for future DIA experiments.

### Quantitative rat organ protein atlas by TMT and DIA-LFQ

We have previously demonstrated that it is possible to baseline resolve the isobaric TMT11-plex report ions in HCD scans recorded with 15,000 resolution and processed with phase-constrained spectrum deconvolution (phi-SDM), which resulted in >50% more identifications with >99% quantified proteins compared to standard settings^23^. This phi-SDM method is implemented as a standard scan option termed Turbo-TMT in the methods editor for the Orbitrap Exploris 480 MS. Consequently, all TMT experiments were recorded with Turbo-TMT to maximize proteome coverage. To fit the 12 organs into a single TMT11-plex experiment, we pooled the duodenum, jejunum and ileum samples into one combined small intestine sample. Likewise, to increase reproducibility in peptide sequencing and enhance quantitative precision by facilitating normalization across the replicate batches, we included a common reference sample created by mixing aliquots of all tissues, in each of the three TMT11-plex batches covering the 10 organs from the three animals (Supplementary Figure 4A). To maximize proteome coverage in the TMT experiments, we performed offline HpH chromatography and collected 46 fractions, which we analyzed by online DDA-based LC-MS/MS using the turbo-TMT method. In contrast, for label-free quantitation we performed single-shot DIA-FAIMS experiments using the individual organ-specific spectral libraries for identification matching in Spectronaut. Consequently, the DIA experiments of the 12 organs analyzed in biological triplicates were done in just 14 hours of LC-MS/MS analysis time, whereas the three TMT11-plex batches required 3 days of total mass spectrometric analysis time (Figure 4A). Both quantification strategies resulted in comparable proteome coverage with more than ten thousand proteins quantified in total. More than eight thousand protein-coding genes were reproducibly quantified in all three TMT experiments, where each TMT set corresponds to one single rat (Supplementary table 4). In contrast, DIA-FAIMS highlights the tissue specificity of the proteomes with variable number of protein groups identified per tissue, from more than 5000 proteins in testis to 2000 proteins on average in muscle (Supplementary figure 4B). Importantly, merging the protein coding genes quantified in each tissue in all three rats resulted in more than 8,000 proteins (Figure 5A and Supplementary table 5), achieving a similar coverage as in TMT. This yields identification rates of more than 600 proteins per minute for DIA (Figure 5B). To remove the well-known batch bias between TMT experiments, we used ComBat normalization^24^ and compared the normalized protein abundance reproducibility between the biological replica experiments for both TMT and DIA. We generally observed high reproducibility analyzing correlations of the log2-transformed protein intensities with R-squared of r^2^=0.93 for DIA (Figure 5C) and r^2^=0.98 for TMT (Figure 5D). The higher correlation for TMT experiments likely reflects the greater number of peptides identified per protein on average as well as the higher precision and ratio compression in TMT compared to label-free quantification^25^. To compare the reproducibility of the three biological replicates of all the rat organ proteomes analyzed by TMT and DIA, respectively, we performed a principal component analysis (PCA) of the normalized protein intensity data for all tissues. Both TMT and DIA data show excellent reproducibility with replica always closer in PCA space than distances to other tissues (Figure 5E and Figure 5F). The component #1 in the DIA based quantification explains more of the variance than for the TMT data - mainly separating the myocyte-rich tissues, cardiac and skeletal muscle, from the other tissues, most likely due to their higher dynamic range of protein abundances. Unsupervised hierarchical clustering of relative protein abundances across tissues reveal very similar protein expression patterns between DIA-FAIMS and TMT experiments, suggesting that both quantification strategies provides very similar biological insights (Figure 5G). A significant advantage in DIA-FAIMS is that it allows us to evaluate the tissue-specificity of the rat proteome. Therefore, we analyzed the distribution of proteins across all tissues to differentiate tissue-specific versus globally expressed proteins. A larger proportion of the identified proteins are tissue specific than commonly expressed with more than two thousand of the identified proteins exclusively found in one tissue and only about one thousand proteins found in all tissues (Figure 5H). By far the majority of tissue-specific proteins were found in testis (Figure 5I), which is in line with observations made in other datasets of global tissue expression profiling^26^. To investigate the biological roles of the tissue-specific versus the globally expressed proteins, we performed a gene ontology (GO) enrichment analysis across the protein-specific tissue distributions, and found that globally expressed proteins are enriched for mitochondrial proteins involved in oxidative phosphorylation, whereas transcription factors^27^ are generally tissue-specific proteins (Figure 5J). These findings are consistent with the fact that cellular housekeeping functions such as ATP production by oxidative phosphorylation is generic to all cell types, whereas tissue specificity is facilitated by temporal and spatial gene expression patterns, which are driven by transcription factors^28^.

### Rapid, sensitive and deep phosphoproteomics

The abundance of a protein is not always a direct measure of its cellular activity, which is often modulated by dynamic post-translational modifications (PTMs)^29^. Site-specific protein phosphorylation is one of the most important PTMs regulating essentially all cellular protein signaling networks by controlling protein activity, subcellular localization, turnover and especially protein-protein interactions^30^. We have previously demonstrated that it is possible to reproducibly identify close to five thousand unique phosphopeptides from 200 μg TiO2-enriched HeLa in-solution digest in just 15 min of LC-MS/MS analysis on an Easy nLC1200 coupled to a Q Exactive HF-X™ instrument^7^. This is equivalent to an analysis throughput of 45 samples per day. We have now streamlined and automated the entire phosphoproteomics workflow on the KingFisher™ Flex robot incorporating on-bead PAC digestion and magnetic Ti-IMAC phosphopeptide-enrichment, which enable parallel preparation of up to 96 phosphopeptide samples (Supplementary Figure 5). Analyzing the phosphopeptides prepared from 200 μg HeLa using data-dependent acquisition with the 28 Hz HCD method on the Orbitrap Exploris 480 MS with the Evosep One using the 60 samples per day method resulted in identification of more than eight thousand unique phosphopeptides of which more than five thousand could be localized confidently^31^ to a single amino acid (Figure 6A). Consequently, this new workflow increases the coverage of the same HeLa phosphoproteome by more than fifty percent in shorter analysis time. To test if this automated phosphoproteomics workflow is also suitable for tissue samples, we analyzed the phosphoproteomes of the 12 organ tissues from the three individual rats. We recently demonstrated that a spectral library-free DIA search, directDIA, of phosphopeptide enriched samples is superior to DDA^32^. We therefore applied this acquisition strategy to analyze the 36 samples, which only required 14 h of MS instrument time, and resulted in the identification of more than 22,000 unique phosphopeptides (Figure 6B). Unsupervised hierarchical clustering of z-scored and log-transformed phosphorylation site intensities revealed high correlation between replica of the same tissue and tissue-specific phosphorylation patterns (Figure 6C). To identify protein kinases that exhibit tissue-enhanced activity, we mapped all known kinase-substrates from PhosphoSitePlus^33^ onto our dataset and calculated the fraction of known substrates identified in our dataset for each kinase in 1 to 12 tissues. This analysis revealed that CAMK2A is the most tissue-specific kinase with one-third of the identified substrates unique to a single tissue (Figure 6D). Cluster analysis highlighted heart tissue as the organ with the highest CAMK2A activity, where it phosphorylates proteins involved in cardiac muscle contraction^34^ (Figure 6E). Conversely, the known substrates of cAMP-dependent protein kinase (PKACA) were more uniformly distributed across the tissues (Figure 6F-G). The fast and sensitive organ phosphoproteomics profiling presented here demonstrated that there are major differences in the phosphorylation patterns across tissues, and accordingly it can provide molecular insights into which and how signaling pathways are regulated in a tissue-specific manner.

**Figure 6.**
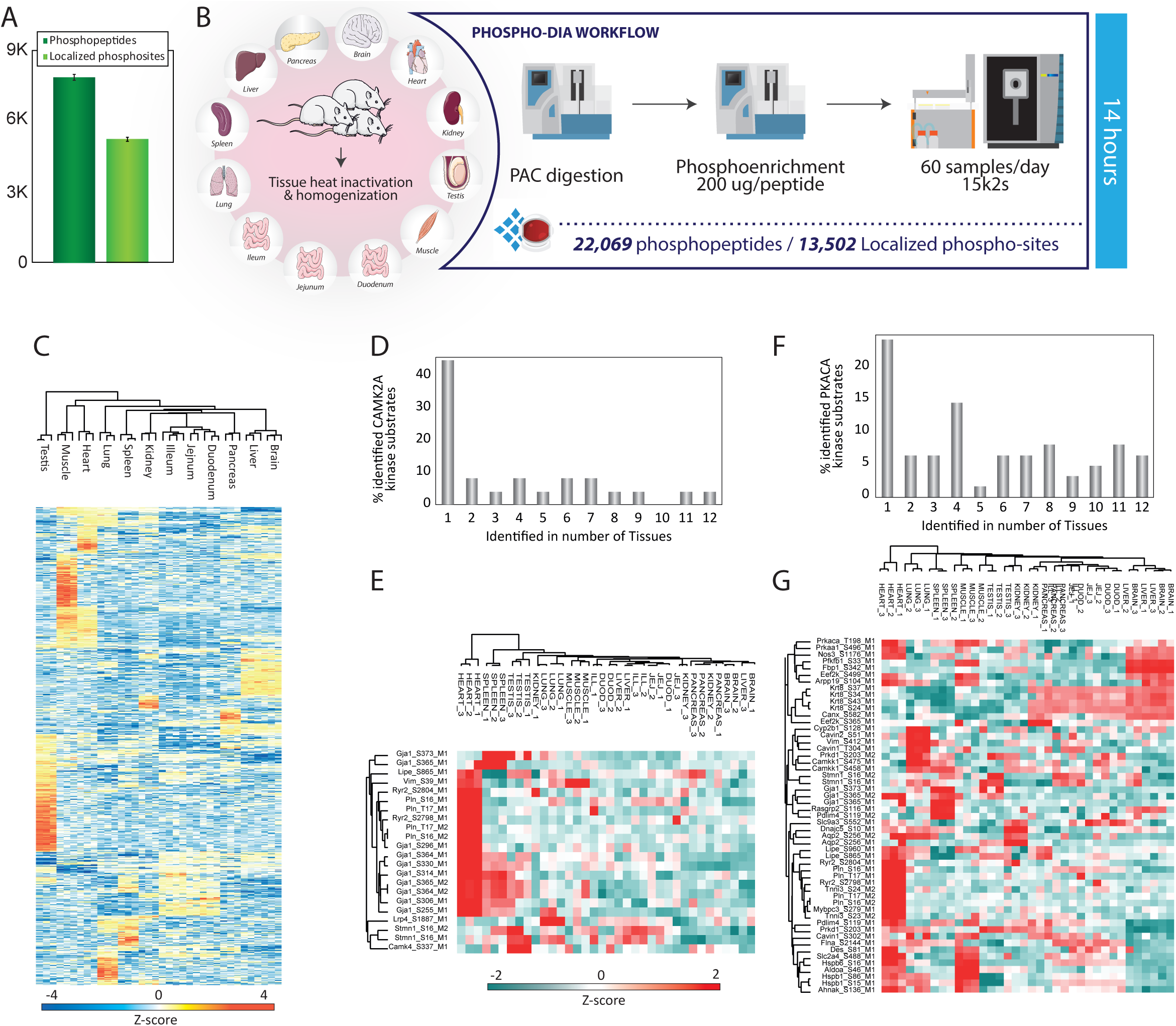
**(a)** Number of phosphopeptides (dark green) and localized phosphosites (light green) from 200 μg of peptides enriched using Ti-IMAC-HP and analyzed with 21 minutes gradient. **(b)** Phospho-DIA workflow of 12 rat tissues from 3 independent rats **(c)** Heatmap of ANOVA regulated phosphorylation sites of the rat tissues depicted as z-score. **(d)** Bar plot showing the percentage of known CAMK2A substrates regulated in different number of tissues. **(e)** Heatmap showing the differential regulation of phosphorylation in CAMK2A substrates across all rat tissues analyzed. **(f)** Bar plot showing the percentage of known PKACA substrates regulated in different number of tissues. **(g)** Heatmap showing the differential regulation of phosphorylation in PKACA substrates across all rat tissues analyzed.

## CONCLUSION

The Orbitrap Exploris 480 MS is a compact hybrid quadrupole-Orbitrap mass spectrometer that provides high-quality HCD tandem mass spectra at acquisition rates of up to 40 Hz, which in combination with the Evosep One system enables routine identification of more than one thousand unique peptides per LC gradient minute. The application of FAIMS with a single CV increased proteome coverage in short gradients by close to fifty percent in data-dependent acquisition based shotgun proteomics experiments of human cells at the expense of lower peptide coverage. Importantly, FAIMS enables analysis of minute sample amounts by effectively removing interfering singly-charged background ions. We demonstrate that it is possible to quantify more than one thousand proteins from less than 50 HeLa cells with DIA-FAIMS, highlighting its potential for applications such as laser microdissected clinical samples and single-cell proteome analysis. The fast sequencing speed, high-sensitivity and robustness of the Orbitrap Exploris 480 MS makes it ideally suited for data-independent acquisition. The combination of FAIMS and data-independent acquisition provides the best of two worlds with more peptides identified and up to 5100 mammalian proteins quantified in single-shot analysis using short LC gradients, allowing up to 60 samples to be measured in 24 hours. This acquisition strategy enables high-throughput and large-scale profiling of hundreds of samples, as demonstrated by the quantitative organ tissue atlas we provide here, which includes a high-quality spectral library consisting of more than two hundred thousand unique peptides covering 12,900 protein-coding genes. This dataset comprises, to our knowledge, the deepest rat proteome to date. This spectral library was recorded by analyzing more than one thousand LC-MS/MS runs of offline high pH reversed-phase peptide fractions back-to-back, which also demonstrates the robustness of the system throughout large-scale applications. In addition to label-free quantification the Orbitrap Exploris 480 MS is also optimized for tandem mass tag based quantification enabled by the fast scanning turboTMT method. Additionally, we have presented a comprehensive comparison between DIA-FAIMS and TMT-based quantification, showing that DIA-FAIMS provides equivalent proteome coverage as isobaric labeling, but without the need for extensive offline fractionation, reducing overall MS analysis time significantly. Interestingly, the observed biological reproducibility in TMT-based quantification was higher than for DIA-FAIMS, but this likely reflects that all tissues from the same animal were combined in one TMT set at the expense of pooling the different parts of the small intestine. This limitation of analyzing only 11 samples within a single TMT experiment is potentially ameliorated with the newly available TMT 16-plex reagents. Finally, the instrument is also a good match for quantitative analysis of post-translational modifications. Sensitive and large-scale analysis of cell line and organ phospho-proteomes is feasible in less than one day of analysis. Consequently, given the high versatility of this instrument, we envision that it will be a workhouse in many proteomics laboratories.

## EXPERIMENTAL PROCEDURES

### Design of a Quadrupole Orbitrap Instrument

The Orbitrap Exploris 480 instrument includes an atmospheric pressure ion source (API) interfaced to a radio-frequency ion funnel via a high-capacity transfer tube, a quadrupole mass filter, a C-trap, an ion routing multipole (IRM), and an ultra-high field Orbitrap mass analyzer. Ions are formed at atmospheric pressure (in this work in a micro-electrospray ion source), pass through the high capacity transfer tube (HCTT) to the ion funnel as described in Martins and coworkers^35^ and then via a mass-selective injection flatapole as described in Scheltema *et al*^15^, into a bent flatapole. The S-shaped bent flatapole is implemented as two parallel printed circuit boards (PCB) with metal rods forming 4 mm gap between its rods, oriented in such a way that the line of sight from the HCTT is open for clusters and droplets to fly unimpeded out of the flatapole onto a dump. Ions are driven through the bent flatapole by an axial electric field created by a voltage divider on PCB stripes. After collisional cooling in the bent flatapole, ions are transmitted via a lens into a hyperbolic segmented-rod quadrupole. A known issue of quadrupole mass filter contamination has been addressed by introducing symmetrical ion loading. In this mode the DC polarity of quadrupole rods is regularly switched so that contamination becomes more symmetrical. After the quadrupole, ions are transferred to the C-trap via a short octapole and a split lens, now integrated with an independent charge detector. Ions fly through the C-trap into a nitrogen filled IRM quadrupole and get trapped or fragmented there. IRM pressure is measured directly by a Pirani gauge and varies in the range from 3 to 10 microbar. As in all other Orbitrap instruments of the Q Exactive^TM7,15,36^ or Orbitrap Fusion™ series^16^, fragmentation of ions in the IRM is achieved by adjusting the offset of the RF rods and the axial field to provide the required collision energy. Similarly, multiple precursor ions could be fragmented at their optimum collision energy without compromising the storage of preceding injections. Unlike previous instruments, the IRM is implemented using the same technology as the bent flatapole, *i.e*. using 4 shaped metal rods soldered to two PCBs separated by precision spacers. Also, a single RF supply drives the C-trap, transport multipole, and IRM at 3.1 MHz. During the return transfer to the C-trap, the same principle of electrodynamic squeezing is applied as in the Orbitrap analyzer. Specifically, IRM offset and C-trap lens voltages are slowly ramped up during the transfer so that ions face an increasing potential barrier as they approach lenses and therefore cannot get deposited on lens edges. Once ions are cooled inside the C-trap, voltages on all its electrodes are raised to the optimum for injection into the Orbitrap analyzer. Unlike all previous designs of C-trap, the pull-out pulse is applied only to the slotted electrode of the C-trap, which ensures strong ion focusing upon extraction so that losses on the edges of the slot are also reduced. While reducing losses and contamination on the exit slit, such extraction requires additional focusing downstream of the C-trap with the help of an additional lens. The symmetrically suspended ultra-high field Orbitrap analyzer is open to pumping on both ends and combines better mechanical and electrical balancing with improved vacuum inside the trap, enabling higher resolution analyzes with acquisition of longer transients. Ions are detected with 4 kV applied to the trap, which is somewhat lower than 5 kV in Q Exactive or Orbitrap Fusion™ series. Nevertheless, this still allows acquisition of transients at resolution settings ranging from 7,500 to 480,000 at m/z 200 (16 and 1024 ms transients, respectively) in 7 steps differing by a factor of 2. The entire ion path from the ion funnel to Orbitrap analyzer is evacuated by a single, purpose-developed, 6-stage turbo pump PM902619A (Pfeiffer Vacuum, Asslar, Germany). UHV is realized without the use of knife-edge sealings by a “chamber-in-chamber” solution in which the pump also serves as a carrier for an inner chamber that houses the C-trap and Orbitrap analyzer. Pressure stages were matched to pumping ports by reducing the length of the hyperbolic-rod quadrupole mass filter, its pre-filters, and transport multipoles, without performance impairment. Ion funnel is evacuated by 120 m3/h single-stage rotary vane pump (Sogevac® SV120 BI FC, Oerlikon Leybold Vacuum, or MS 120, Agilent Technologies) and turbopump is backed by a dedicated Sogevac SV15 pump.

### Processing of Transients

Transients detected in the Orbitrap mass analyzer are processed using an enhanced version of Fourier Transformation (eFT™) as described in Lange *et al*^37^. For TMT experiments, narrow m/z regions near TMT-10 reporter peaks^38,39^ were processed by Phase-constrained signal deconvolution method (Phi-Transform) as described in ^23^. This allowed use of special transient durations of 32 ms (resolution setting 15,000 at m/z 200) specifically for resolving isobaric TMT reporter ions in TMT 11-plex experiments.

### FAIMS Pro

As the front-end of the new mass spectrometer is essentially identical to that of Orbitrap Fusion Lumos™ instrument, the same FAIMS Pro™ option could be used as in the latter. Details of FAIMS Pro design and its characterization are presented in Pfammatter *et al*^13^. The unique feature of FAIMS Pro is the use of cylindrical electrodes that help to focus ions through the electrode assembly as they get carried by nitrogen carrier gas from front to back. With inner electrode blocking “line of sight” for clusters and dust particles, the additional benefit of interface protection from environmental effects is achieved. As this work utilizes higher flow rates than in previous nanospray experiments, method of FAIMS Pro operation includes an additional step of increased FAIMS carrier gas setting at the beginning of the LC gradient.

### Cells

Human epithelial cervix carcinoma HeLa cells were cultured in DMEM (Gibco, Invitrogen), supplemented with 10% fetal bovine serum, 100U/mL penicillin (Invitrogen), 100μg/mL streptomycin (Invitrogen), at 37°C, in a humidified incubator with 5% CO2.

### Lysis and Digestion

Cells were harvested at ∼80% confluence by washing twice with PBS (Gibco, Life technologies) and subsequently adding boiling lysis buffer^16^ (5% sodium dodecyl sulfate (SDS), 5 mM tris(2-carboxyethyl)phosphine (TCEP), 10 mM chloroacetamide (CAA), 100 mM Tris, pH 8.5) directly to the plate. The cell lysate was collected by scraping the plate and boiled for an additional 10 minutes followed by micro tip probe sonication (Vibra-Cell VCX130, Sonics, Newtown, CT, USA) for 2 minutes with pulses of 1 s on and 1 s off at 50% amplitude. Protein concentration was estimated by BCA.

### Rat tissue preparation

The study was carried out following approved national regulations in Denmark and with an animal experimental license granted by the Animal Experiments Inspectorate, Ministry of Justice, Denmark. Three male Sprague Dawley rats (Crl:SD, male, 200 g, Charles River, Germany) were anesthetized with isoflurane. The animals were perfused (1.5 min, 30 mL/min) with isotonic saline containing protease inhibitors (0.120 mM EDTA, 14 μM aprotinin, 0.3 nM valine-pyrrolidide and Roche Complete Protease Inhibitor tablets (Roche), pH = 7.4). Tissues were quickly removed and snap frozen in isopentane on dry ice. Frozen organs were heat inactivated (Denator T1 Heat Stabilizor, Denator) and a portion of each tissue was transferred to an SDS solution (5% SDS, 25 mM Tris, pH 8.5) and homogenized by ceramic beads using steps of 20 s at 5500 rpm (Precellys 24, Bertin Technologies) until all tissue was clarified. The tissues were heated for 10 min at 95°C followed by micro tip sonication on ice and clarified by centrifugation (20 min, 16,000g, 4°C). Samples were reduced and alkylated by adding 5 mM tris(2-carboxyethyl)phosphine and 10 mM chloroacetamide for 20 min at room temperature.

### Automated PAC digestion

Protein digestion was automated on a KingFisher™ Flex robot (Thermo Scientific) in 96-well format. The 96-well comb is stored in plate #1, the sample in plate #2 in a final concentration of 70% acetonitrile and with magnetic Amine beads (ReSyn Biosciences) in a protein:bead ratio of 1:2. Washing solutions are in plates #3-5 (95% Acetonitrile (ACN)) and plates #6-7 (70% Ethanol). Plate #8 contains 300 μl digestion solution of 50 mM ammonium bicarbonate (ABC), LysC (Wako) in an enzyme/protein ratio of 1:500 (w/w) and trypsin (Sigma Aldrich) in an enzyme:protein ratio of 1:250. The protein aggregation was carried out in two steps of 1 min mixing at medium mixing speed, followed by a 10 min pause each. The sequential washes were performed in 2.5 min and slow speed, without releasing the beads from the magnet. The digestion was set to 12 hours at 37 degrees with slow speed. The workflow is illustrated in Supplementary Figure 5A. Protease activity was quenched by acidification with trifluoroacetic acid (TFA) to a final concentration of 1%, and the resulting peptide mixture was concentrated on Sep-Pak (C18 Classic Cartridge, Waters, Milford, MA, USA). Peptides were eluted with 2 mL 40% ACN, followed by 2 mL 60% ACN. The combined eluate was reduced by SpeedVac (Eppendorf, Germany) and the final peptide concentration was estimated by measuring absorbance at 280 nm on a NanoDrop 2000C spectrophotometer (Thermo Fisher Scientific). Peptides were preserved at -80 degrees for further analysis or loaded on EvoTips according to the manufacturer’s protocol.

### High-pH Fractionation

Peptides were fractionated using a reversed-phase Acquity CSH C18 1.7 μm 1 × 150 mm column (Waters, Milford, MA, USA) on an UltiMate 3000 high-pressure liquid chromatography (HPLC) system (Thermo Fisher Scientific) operating at 30 μL/minute. Buffer A (5 mM ABC) and buffer B (100% ACN) were used. Peptides were separated by a linear gradient from 5% B to 35% B in 55 minutes followed by a linear increase to 70% B in 8 minutes. 46 fractions were collected without concatenation.

### TMT Labeling

Rat peptides were TMT-labeled according to the overview in Supplementary Figure 4A with 10 μg of peptide and 4 μl of TMT reagent per channel. Labeling conditions were performed according to manufacturer’s protocol. After labeling samples were pooled, acidified and concentrated on SepPaks (C18 Vac C18 Cartridge, 1cc/50mg 55-105 um Waters, Milford, MA, USA). Peptides were eluted with 300 μl 40% ACN, followed by 300 μl 60% ACN and by 300 μl 80% ACN. The combined eluate was reduced by SpeedVac, resuspended in 5 mM ABC and offline high pH reversed phase fractionated as described above.

### Automated Phosphopeptide Enrichment

Phosphopeptide enrichment was carried out on a KingFisher™ Flex robot (Thermo Fisher Scientific) in 96-well format as previously described (Ref) and also outlined in Supplementary Figure 5B. 200 μg of peptide were used for enrichments with 30 μl of magnetic Ti-IMAC HP beads (a prototype phosphopeptide enrichment product from ReSyn Biosciences), which is an improved version of the commercially available magnetic Ti-IMAC beads (ReSyn Biosciences). Briefly, the 96-well comb is stored in plate #1, 30 μl Ti-IMAC HP beads in 100% ACN in plate #2 and loading buffer (1 M glycolic acid, 80% ACN, 5% TFA) in plate #3. The sample volume is minimum doubled with loading buffer while kept in a total of 300 μl and added in plate #4. Plates 5-7 are filled with 500 μl washing solutions; loading buffer, 80% ACN, 5% TFA and 10% ACN, 0.2% TFA respectively. Plate #8 contains 200 μl 1% ammonia for elution. The beads were washed in loading buffer for 5 min at medium mixing speed, followed by binding of the phosphopeptides for 20 min and medium speed. The sequential washes were performed in 2 min and fast speed. Phosphopeptides were eluted in 10 min at medium mixing speed. The eluate is acidified with TFA and loaded directly on EvoTips.

### LC–MS/MS

All samples were analyzed on the Evosep One system using an in-house packed 15 cm, 150 μm i.d. capillary column with 1.9 μm Reprosil-Pur C18 beads (Dr. Maisch, Ammerbuch, Germany) using the pre-programmed gradients (60, 100 and 200 samples per day). The column temperature was maintained at 60°C using an integrated column oven (PRSO-V1, Sonation, Biberach, Germany) and interfaced online with the Orbitrap Exploris 480 MS. Spray voltage were set to 2 kV, funnel RF level at 40, and heated capillary temperature at 275°C. For DDA experiments full MS resolutions were set to 60,000 at m/z 200 and full MS AGC target was 300% with an IT of 25 ms. Mass range was set to 350-1400. AGC target value for fragment spectra was set at 200% with a resolution of 15,000 and injection times of 22 ms and Top12. Intensity threshold was kept at 2E5. Isolation width was set at 1.3 m/z. Normalized collision energy was set at 30%. For DIA experiments full MS resolutions were set to 120,000 at m/z 200 and full MS AGC target was 30% with an IT of 45 ms. Mass range was set to 350-1400. AGC target value for fragment spectra was set at 100%. 49 windows of 13.7 Da were used with an overlap of 1 Da. Resolution was set to 15,000 and IT to 22 ms. Normalized collision energy was set at 27%. All data were acquired in profile mode using positive polarity and peptide match was set to off, and isotope exclusion was on. Default settings were used for FAIMS with voltages applied as described in the manuscript, except gas flow, which was applied with 3 L/min in the first minute of the method and spray voltage which was set to 2.3 kV. For TMT experiments the turbo-TMT was enabled, MS/MS data was recorded in centroid mode. Isolation window was set to 0.7 Da and first mass was set to 100 m/z.

### Rat tissue specific library generation

A pool from each tissue was generated using 40 μg of peptide from a given tissue from each independent rat. Each of the tissue pools were fractionated using high-pH reversed phase fractionated as described above into 46 fractions. For each tissue, the fractionated proteome was analyzed twice on the Evosep One with the column details described above and using the pre-programmed gradient (200 samples per day). Around 250 ng from each fraction was analyzed on a Q-Exative HF-X™ using a MS/MS resolution of 15,000, injection time of 22 ms and a Top12 method. Another 250 ng was then analyzed on an Orbitrap Fusion Lumos with FAIMS using a MS/MS resolution of 30,000, injection time of 54 ms and a Top6 method and a CV of -45 V for FAIMS.

### Raw Data Processing

All Data-dependent analysis (DDA) raw files were processed with MaxQuant^19^ v1.6.7.0 using the integrated Andromeda Search engine^40^. All HeLa data is searched against the human Uniprot Reference Proteome without isoforms, while all rat files are searched against the rat Uniprot Reference Proteomes with and without isoforms and the mouse Uniprot Reference Proteome without isoforms (all with January 2019 release). Trypsin was specified as enzyme, cleaving after all lysine and arginine residues and allowing up to two missed cleavages. Carbamidomethylation of cysteine was specified as fixed modification and protein N-terminal acetylation, oxidation of methionine, deamidation of Asparagine and Glutamine and pyro-glutamate formation from glutamine were considered variable modifications with a total of 2 variable modifications per peptide. “Maximum peptide mass” were set to 7500 Da, the “modified peptide minimum score” and “unmodified peptide minimum score” were set to 25 and everything else was set to the default values, including the false discovery rate limit of 1% on both the peptide and protein levels. The TMT proteomes were searched independently with further specification of TMT 11-plex as label with a reporter ion mass accuracy of 0.002 Da. For the rat proteomes, 12 different searches were performed independently with the “match between runs” feature enabled between fractions. All Data-independent analysis (DIA) raw files were processed with Spectronaut version 13 (Biognosys, Zurich, Switzerland). Project specific spectral libraries were imported from the separate MaxQuant analyzes of the combined analysis of the 2x 46 pre-fractionated fractions of each tissue, and DIA files were analyzed using default settings disabling the PTM localization filter. Phospho-enriched samples from rat tissues were processed in Spectronaut using a library-free approach (directDIA) including phosphorylation on STY as a variable modification.

### Data analysis

DIA results from each independent tissue were merged by gene name. For quantification in DIA, we used standard settings in Spectronaut for quantification at MS2 level. Proteins, which were not quantified in all three replicates for at least one tissue were discarded for further analysis. Missing values were imputed in Perseus using values from a normal distribution (width=0.3, downshift=1.8). For quantification in TMT analysis, Reporter intensities from proteinGroups.txt table were used. TurboTMT data was filtered to remove proteins not quantified in all channels in three independent experiments. Intensities across channels were log2 transformed and normalized using quantile normalization. Batch effect between independent TMT experiments was removed using ‘ComBat’ function implemented in the sva package^24^ in R. For clustering, TMT and DIA data were separately transformed by z-score. For comparison with TMT, the average of duodenum, jejunum and ileum values for each independent rat was used. Finally, proteins were clustered using Euclidean distance in Perseus. Principal Component Analysis (PCA) was performed using proteins quantified in all experiments (either DIA or TMT), using the ‘prcomp’ function in R. Phospho-DIA data was searched with directDIA in Spectronaut and the report output collapsed to either “modification-specific-peptide” or “phospho-site” tables using the Perseus plugin previously described^32^. Phosphosite intensities were log2 transformed and missing values were imputed in Perseus using values from a normal distribution (width=0.3, downshift=1.8). Afterwards, intensities were normalized using loess function implemented in the limma package in R, and ANOVA test was performed using an FDR < 1%. Phosphosite intensities were z-score transformed and clustered using Euclidean distance.

## AUTHOR INFORMATION

### Notes

The authors declare the following competing financial interest(s): K.F., T.N.A., A.H. and A.M. are employees of Thermo Fisher Scientific, the manufacturer of the Orbitrap Exploris 480 MS instrument used in this research.

## ACKNOWLEDGMENTS

Part of this work has been funded as part of the MSmed project that has received funding from the European Union’s Horizon 2020 Research and Innovation program under grant agreement no. 686547. We thank our colleagues at Thermo Fisher Scientific, especially Jan-Peter Hauschild, Amelia Peterson, Erik Couzijn, Eduard Denisov, Denis Chernyshev, Christian Hock, Hamish Stewart, Ralf Hartmer, Christian Thoeing, Oliver Lange, Mathias Mueller, Arne Kreutzmann, Wilko Balschun, Aivaras Venckus, Alexander Kholomeev, Gregor Quiring, Frank Czemper, Andreas Wieghaus, Michael Belford, Julia Kraegenbring, Alexander Harder, Kerstin Strupat, Markus Kellmann. Work at The Novo Nordisk Foundation Center for Protein Research (CPR) is funded in part by a generous donation from the Novo Nordisk Foundation (NNF14CC0001). JVO and PR are supported by the Marie Skłodowska Curie European Training Network “TEMPERA” (grant number 722606). The authors would like to thank Stoyan Stoychev and Justin Jordaan for great help and input for optimizing our phospho-enrichment workflow and establishing automated platform for sample preparation. We would also like to thank Nicolai Bache for help and input on the chromatographic methods as well as Anna Secher for providing rat tissues and all members of the Olsen Group for critical input on the manuscript.

## ABBREVIATIONS

ACN: acetonitrile;
APD,DDA: data dependent acquisition;
DIA: data independent acquisition;
FAIMS: high-field asymmetric waveform ion mobility spectrometry;
FA: formic acid;
HCD: higher-energy collisional dissociation;
HCTT: high capacity transfer tube;
IT: maximum injection time;
LC: liquid chromatography;
m/z: mass-to-charge;
MS: mass spectrometry;
MS/MS: tandem mass spectrometry;
PSMs: peptide spectrum matches;
PTMs: post translational modifications;
TFA: trifluoroacetic acid;
TMT: tandem mass tags.

## FIGURE LEGENDS

**Supplementary Figure 1.**
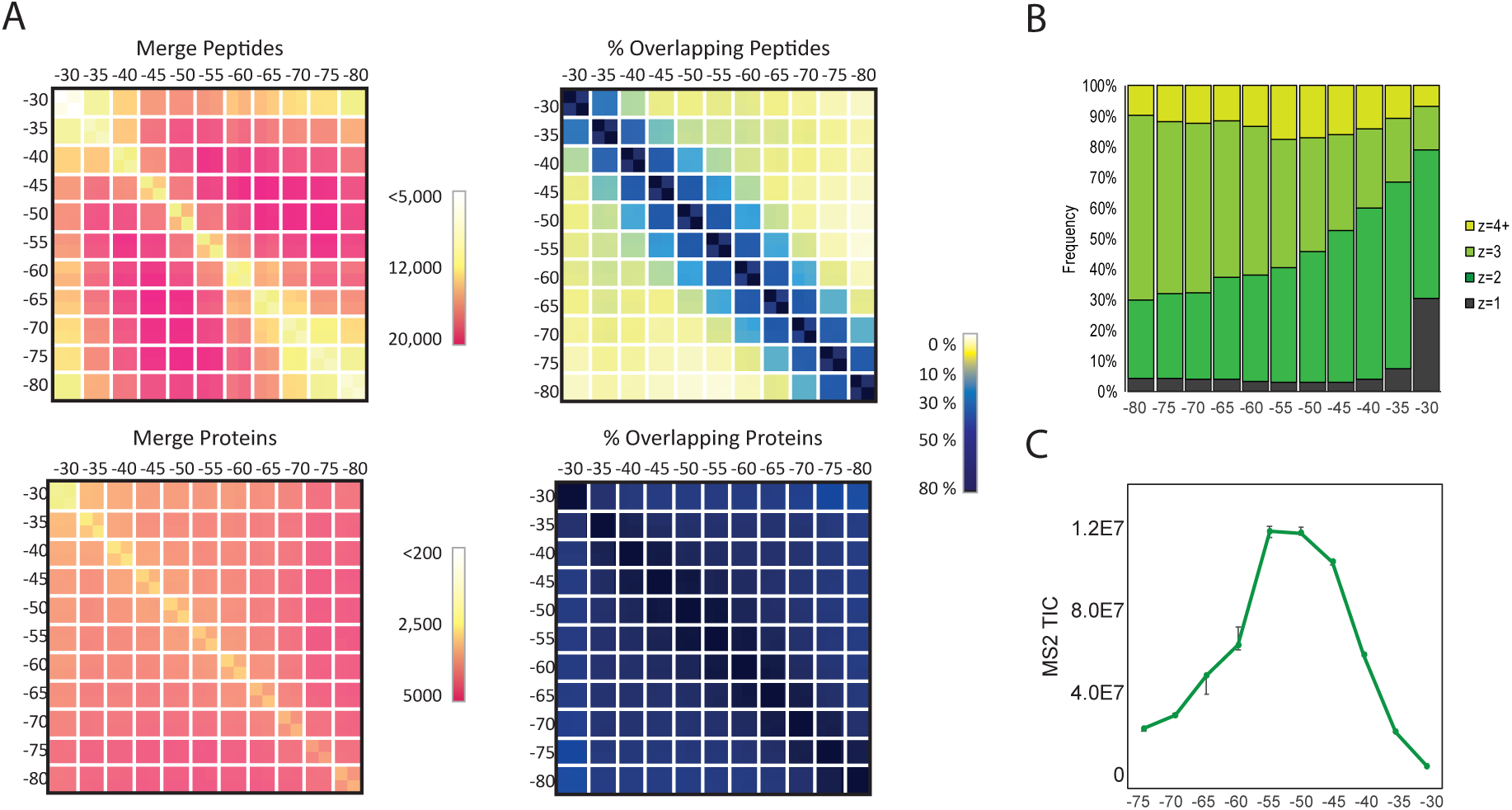
**(a)** Heatmap showing the number of peptides or proteins gained by merging independent DDA-FAIMS runs with different compensation voltages. On the right, heatmap showing the percentage of common peptides or proteins identified in independent DDA-FAIMS runs with different compensation voltages. **(b)** Precursor charge distribution using different compensation voltages. **(c)** MS2-TIC average values at different compensation voltages from 500 ng peptide in a 21 minutes gradient using DDA settings.

**Supplementary Figure 2.**
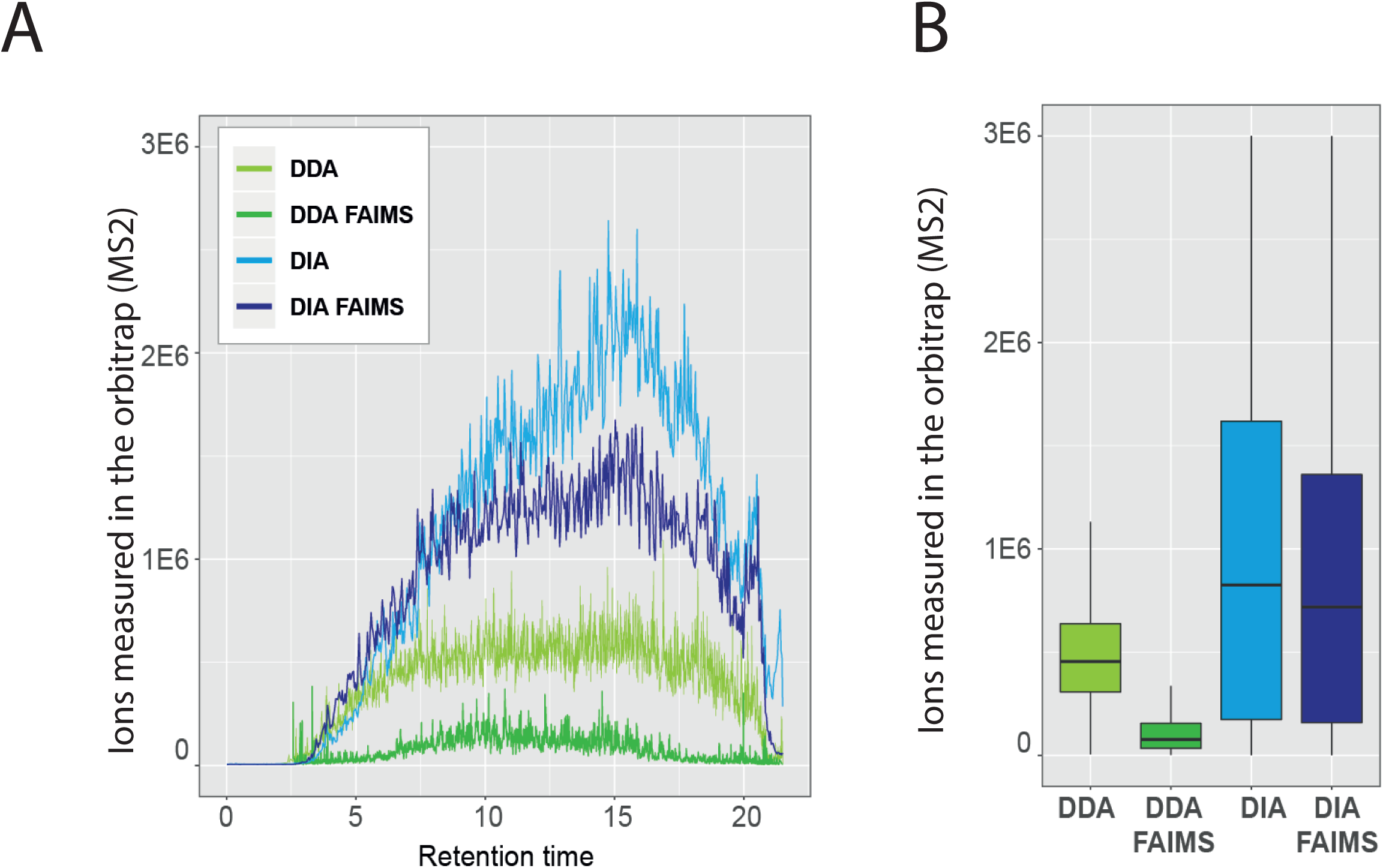
**(a)** Comparison of MS2 ion intensity levels across time in DDA, DDA-FAIMS, DIA and DIA-FAIMS runs using 500 ng peptide. **(b)** Boxplot showing the MS2 ion intensity distribution in DDA, DDA-FAIMS, DIA and DIA-FAIMS runs using 500 ng peptide.

**Supplementary figure 3.**
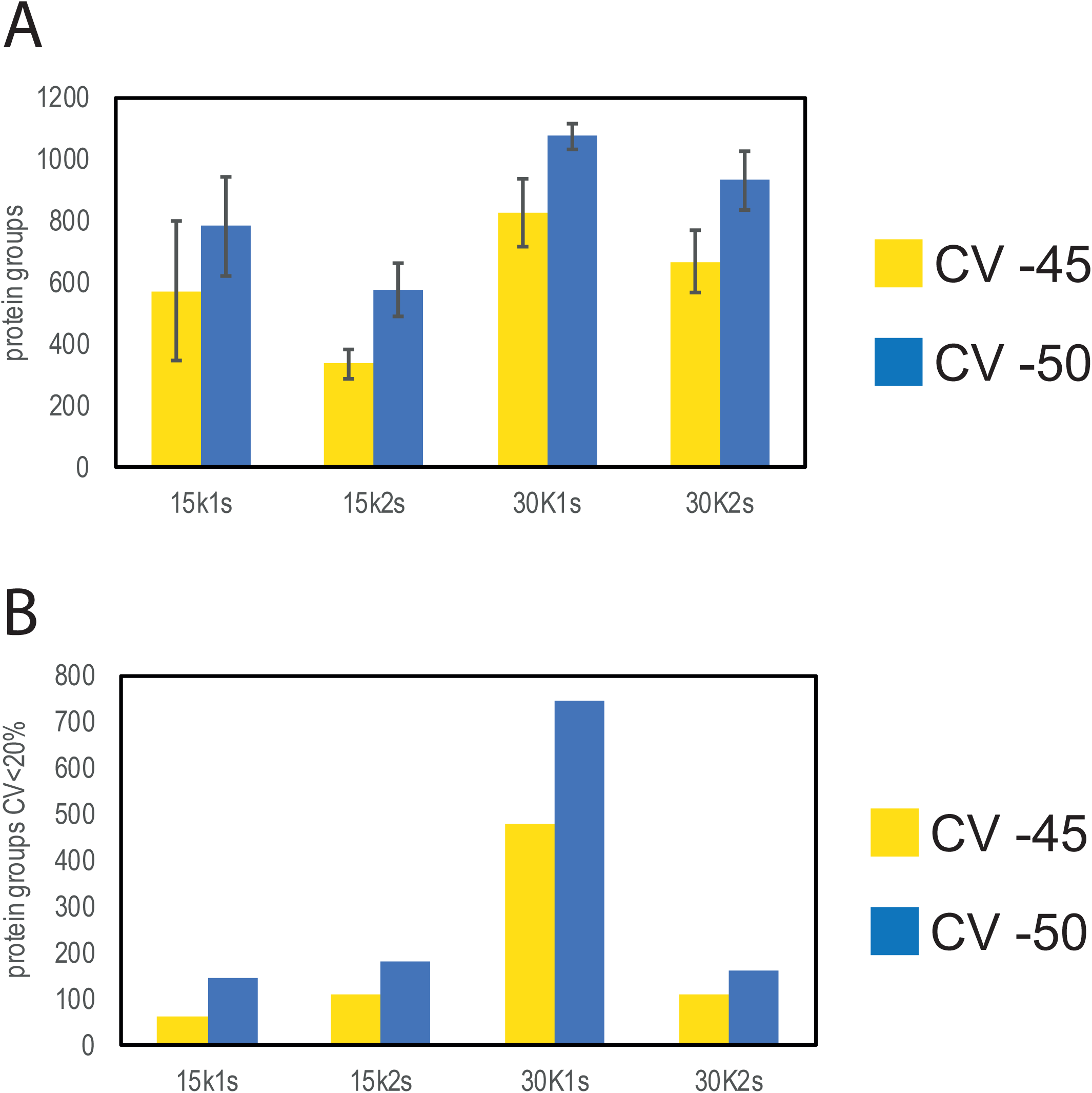
**(a)** Protein groups identified in total using different acquisition methods, cycle times and CV in DIA-FAIMS mode using 5 ng of peptide in a 5 minute gradient (200 samples per day). **(b)** Proteins with a coefficient of variation below 20% for the same parameters as in 3a.

**Supplementary figure 4.**
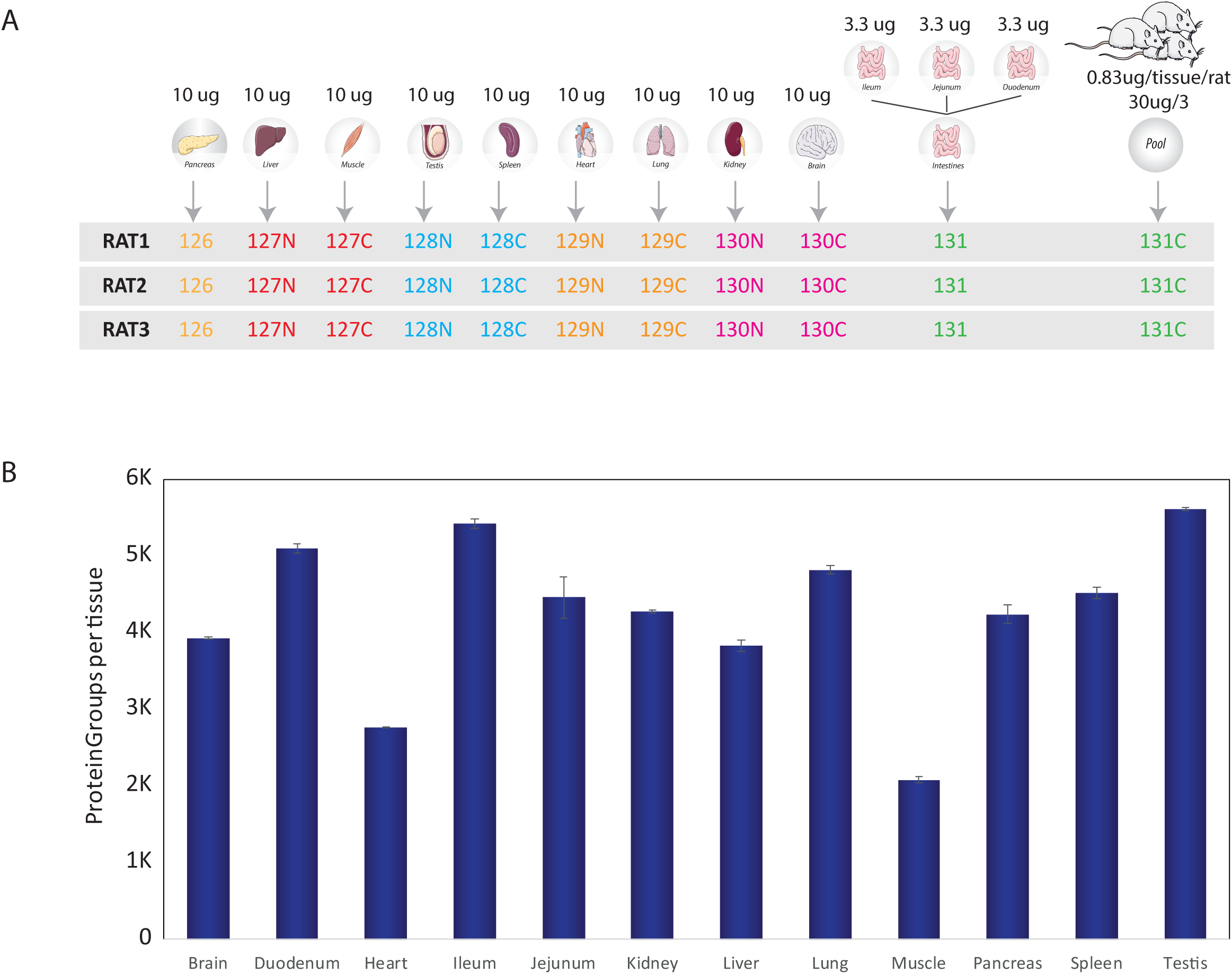
**(a)** TMT labeling scheme for full proteome analysis. **(b)** Bar chart showing the number of protein groups per tissue quantified in each independent tissue in DIA-FAIMS analysis. Plotted value is the average of three replicates, and the error bars represent the standard deviation among them.

**Supplementary figure 5.**
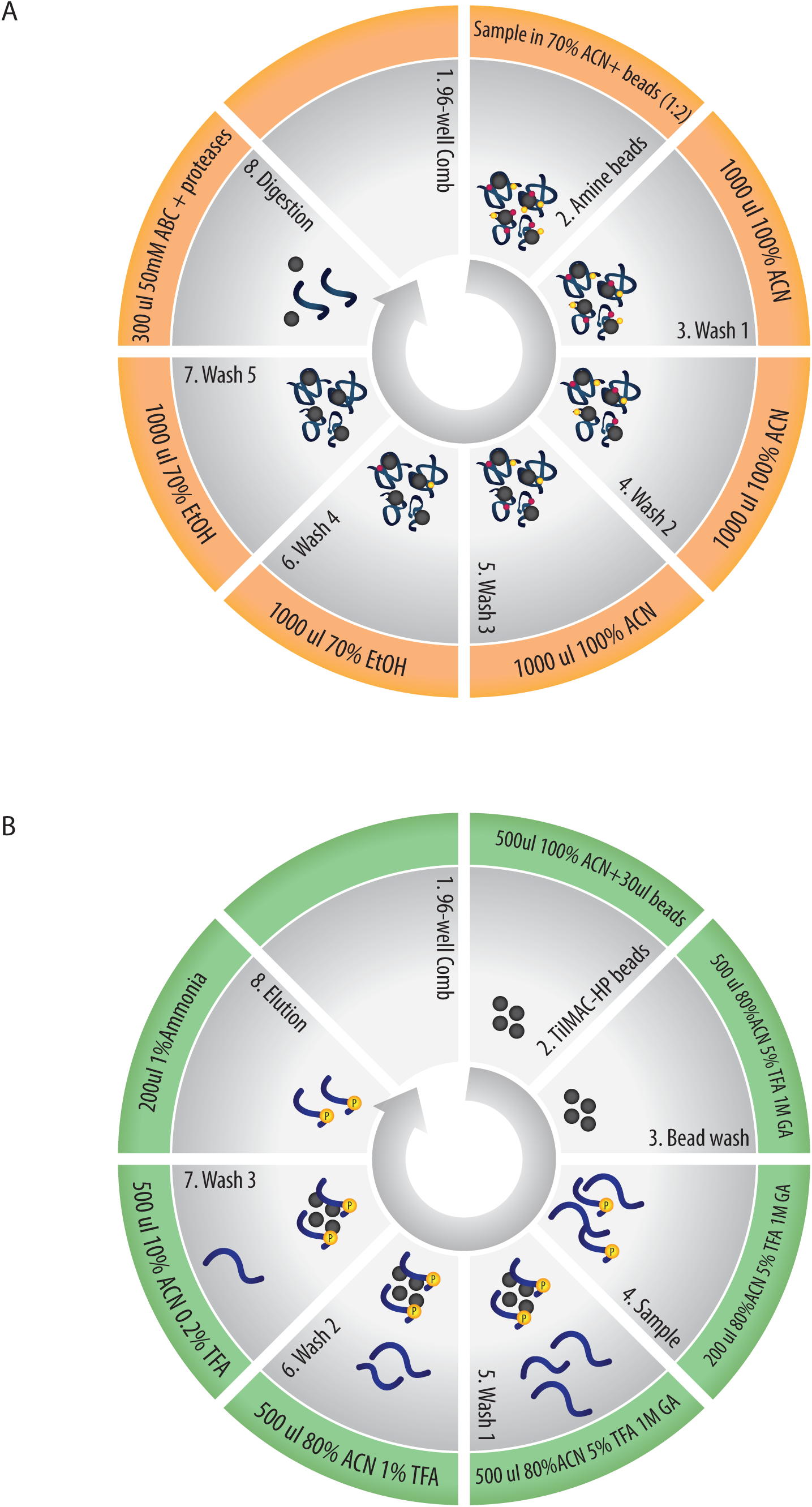
**(a)** Detailed PAC digestion workflow on the KingFisher Flex robot. **(b)** Detailed phospho-enrichment workflow on the KingFisher Flex robot.

